# Tropical Peatlands as Hotspots of Hidden Arcellinida Diversity: Insights from Maludam National Park, Borneo (Sarawak, Malaysia)

**DOI:** 10.64898/2026.10.30.763046

**Authors:** Ángel García-Bodelón, Petr Heděnec, Sharon Yu Ling Lau, María Martínez-Ríos, Umar Hussaini Tarmizi, Meldon Fienberg Kelbin, Kim San Lo, Laurina Lenny Kambit, Lulie Melling, Nura ElKhouri-Vidarte, Carmen Soler-Zamora, Enrique Lara

**Author notes:** Sarawak Research and Development Council, Lot 6035, Kuching-Samarahan Expressway, 94300 Kota Samarahan, Sarawak, Malaysia (Sharon Yu Ling Lau, Meldon Fienberg Kelbin, Kim San Lo, Laurina Lenny Kambit). UN Sustainable Development Solutions Network, Asia Headquarters, Sunway University, 47500 Bandar Sunway, Selangor, Malaysia. Corresponding author: Ángel García-Bodelón, Address: Real Jardín Botánico (RJB-CSIC), C/ Moyano 1, 28014 Madrid, Spain.

## Abstract

Recent investigations on tropical peatlands have shown their extremely high efficiency as carbon sinks, far beyond better studied boreal bogs. The microbial diversity found in those ecosystems, as well as their corresponding ecological and metabolic processes, are just starting to be revealed, and differ considerably from high latitudes systems. However, these efforts have been concentrated almost exclusively towards prokaryotes and fungi, leaving protist communities virtually unknown. Arcellinida (testate amoebae) are well acknowledged to play an important role in boreal and temperate peatland ecology by driving microbial community composition as top predators, but also as nutrient recyclers and carbon fixers. Yet, to our knowledge, no community-level study has addressed their diversity and its drivers in tropical peatlands. In this study, we surveyed Arcellinida communities’ diversity across five functionally distinct vegetation–hydrological zones situated along a hydrological transect in a coastal tropical peatland in the Maludam National Park, Malaysia. Taxonomic composition of Arcellinida communities, characterized by a high diversity of *Trigonopyxis*-related taxa, indicated fundamental differences in diversity and community composition from extratropical ecosystems. Species richness was significantly higher in the zones with developed understory vegetation. Furthermore, the composition of diversity varied significantly among vegetation-hydrological zones. Altogether, this study shows unique Arcellinida communities in a Southeastern Asian peatland and suggests the importance of vegetation-hydrological characteristics for protist diversity.

## INTRODUCTION

Tropical peat swamp forests, or tropical peatlands for short represent unique and highly specialized ecosystems found in equatorial latitudes in South America, Africa and Southern and Southeast Asia. They constitute a unique dual ecosystem, combining the characteristics of both peatland and rainforest (Melling, 2016); they occur on water logged soils and accumulate plant matter at a higher rate than it decomposes, a general characteristic of peatlands that designate them as carbon sinks (IPBES, 2019). High temperatures and humidity drive fast turnover, and carbon accumulation happen faster than in the better studied high latitude peatlands (Hodgkins et al., 2018). These biomes are home to fragile and highly specific organisms such as the fish *Paedocypris progenetica* Kottelat, Britz, Tan & Witte, 2006, the smallest known vertebrate in the world (Kottelat et al., 2006), or the frog *Microhyla borneensis* Parker, 1928, associated with microhabitats provided by the endemic carnivorous plant *Nepenthes ampullaria* Jack, 1835 (Das & Haas, 2010). In plants, stressful hydrological conditions have also driven structural and physiological adaptations for oxygen acquisition in waterlogged soils, for instance, developing pneumatophores (breathing roots) in peat swamp trees (Busman, Melling, et al., 2023).

Despite above-ground diversity in tropical peatland is starting to be well catalogued, the below-ground communities that drive main ecological processes such as decomposition and transformation of organic matter in tropical peat soils remain unexplored. Research on decomposer groups (i.e. bacteria and fungi) in South Eastern Asia is revealing a wealth of unreported organisms and suggests unreported biochemical processes (Birnbaum et al., 2025; Morrison et al., 2021a). Protists, the main regulators of below-ground decomposer communities (Creevy et al., 2018) are virtually unexplored in tropical peatlands. Studying all protist groups together implies considering organisms that differ deeply in their size, life stage and ecology (Geisen et al. 2018). Instead, Arcellinida (lobose testate amoebae) are possibly the best studied microbial group in boreal and temperate peatlands. Collectively, they can represent half of the *Sphagnum-*associated microbial biomass in pristine peatbogs (Gilbert et al., 1998). As top predators, they regulate populations of other heterotrophic microbial predators (including small animals), algae and fungi (Gilbert et al., 2003) and play therefore an essential role in nutrient cycling. Moreover, the symbiotic association of certain species with trebouxiophycean algae gives them a non-negligible role in carbon fixing at the whole ecosystem level, about 13% (Jassey et al., 2015). Besides their active ecological role, testate amoebae communities are excellent bioindicators that provide information on peatland hydrological conditions (Booth, 2008) or nutrient status (Lamentowicz et al., 2013).

In the last years, a few studies have been exploring Arcellinida diversity in tropical peatlands in the Amazon (Swindles et al., 2014), the Panamanian coast (Swindles et al., 2018a) and Central Sumatra (Krashevska et al., 2020). All these works showed a relatively modest diversity, composed mostly of cosmopolitan morphotypes. A common observation was a succession of communities all along water table depth gradients and pH; sea level influence appeared to play also a role in species distribution (Swindles et al., 2018b). However, all these studies were based on the microscopic observations of shells, which is known to underestimate diversity as compared to metabarcoding approaches, thus blurring fine distribution patterns (González-Miguéns et al., 2023).

Potential differences in organism diversity will be influenced by the intrinsic and functional characteristics of the peatlands in question. Like their temperate and boreal counterparts, tropical peatlands are characterized by high precipitation, acidic and waterlogged anoxic conditions, and store carbon as non-decomposed plant material (Busman, Maie, et al., 2023; Melling, 2016). But in contrast to boreal and temperate peatlands, which are largely dominated by bryophytes (like *Sphagnum* spp.) and other non-woody vegetation, tropical peat substrate derives mostly from woody forest material, including roots, trunks, branches and leaves (Page et al., 1999, 2006). The peat-building process changes also; in boreal and temperate peatlands it is primarily controlled by low temperatures while in the tropics it is shaped by high temperatures and very intensive rainfall exceeding 2000 mm annually (Andriesse, 1988; Zinck, 2011). While both ecosystem types are widely recognized as carbon sinks, tropical peatlands generally exhibit higher long-term carbon accumulation rates than boreal and temperate peatlands owing to greater primary productivity; nevertheless, high latitude peatlands store a substantially larger proportion of the global peat carbon pool because of their much greater areal extent (Dommain et al., 2011; K. Ribeiro et al., 2021; Yu, 2012). However, fundamental differences exist with boreal and temperate peatlands; the microbiome of boreal and temperate peatlands has a strong potential for plant material degradation but lacks enzymes for complete denitrification. However, tropical peatlands microbiomes have a broad potential for all key peatland biogeochemical processes across all depths (Cheung et al., 2025). These different functions hold a great potential for new taxa and consortia discovery (Morrison et al., 2021b).

In this work, we surveyed the diversity of Arcellinida in a coastal tropical peatland using a group-specific metabarcoding approach, to our knowledge for the first time. Our study site is located in a natural dome-shaped coastal peatland with minimal anthropogenic perturbations within the Maludam National Park in Sarawak (Malaysia). We sampled through a transect spanning from the upper tidal zone to the upper part of the peat dome, running across a water table gradient and five vegetation-hydrological zones. Our aims are therefore to (1) provide the first molecular taxonomic characterization of Arcellinida communities in a tropical peatland, with particular emphasis on their comparison with temperate counterparts; and (2) assess the extent to which these communities vary among vegetation–hydrological zones, given that these habitats are known to host highly distinct prokaryotic communities (Dom et al., 2021).

## MATERIALS AND METHODS

### Study Area and Field Sampling

The Maludam peatland is located in the Maludam National Park in Sarawak (Malaysia), at the shore of the Batang Lupar estuary. This particular peatland forms a convex coastal peat dome in which surface elevation and peat thickness generally increase from the estuarine margin towards the interior. Peat thickness reaches approximately 12 m near the center of the dome (Melling, 2016). This topographic gradient is accompanied by changes in tidal influence, water-table position, peat characteristics, nutrient availability and vegetation composition.

Across this peatland, five vegetation–hydrological zones were selected for sampling along a transect extending from the Batang Lupar estuary towards the interior of the peat dome (Fig. 1). Following Melling (2016), these zones were classified as:

**Figure 1.**
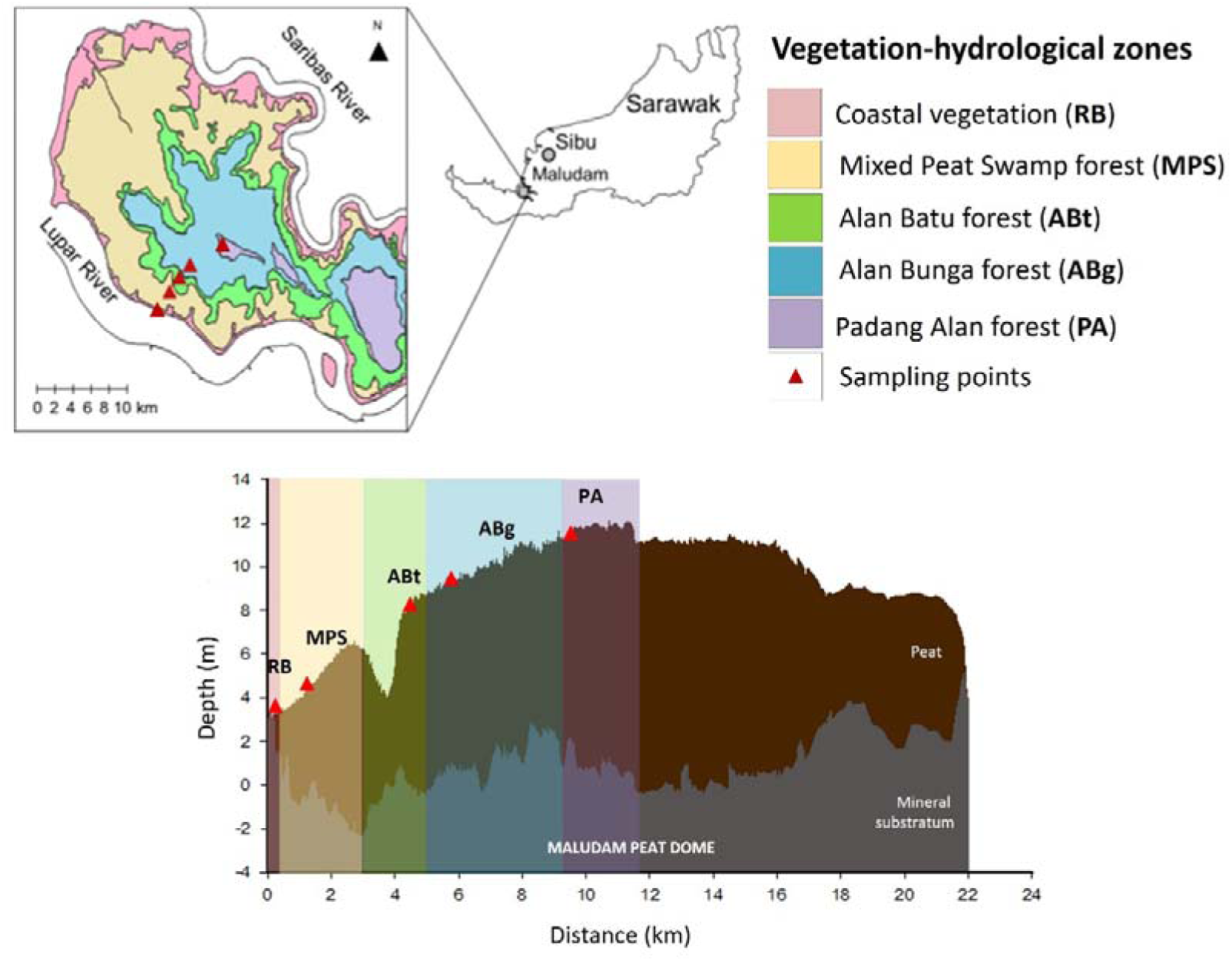
Map of Maludam National Park showing the distribution of vegetation-hydrological zones and sampling locations. There are five vegetation-hydrological zones: Coastal vegetation (RB), Mixed Peat Swamp (MPS), Alan Batu forest (Abt), Alan Bunga forest (ABg), and Padang Alan forest (PA). The elevation profile along the transect graph with coloured bands indicating the corresponding vegetation-hydrological zones and red triangles marking the sampling locations. Adapted from Dom et al. (2021).

#### Coastal vegetation (river bank)

A tidally influenced zone occurring near the estuary and characterized by *Nypa fruticans*, *Rhizophora* spp., *Sonneratia* spp. and other mangrove-associated vegetation.

#### Mixed Peat Swamp forest

A structurally complex and species-rich forest with an uneven canopy, abundant climbers and epiphytes, and relatively sparse understory vegetation. Characteristic tree species include *Gonystylus bancanus*, *Dactylocladus stenostachys* and *Copaifera palustris*.

#### Alan Batu forest

A peat swamp forest structurally similar to the mixed peat swamp forest but containing scattered large individuals of *Shorea albida*. *Stemonurus umbellatus* is also characteristic of this zone.

#### Alan Bunga forest

A forest strongly dominated by large *Shorea albida* trees, with an even upper canopy reaching approximately 45–60 m and a relatively dense understory. Climbers and epiphytes are uncommon.

#### Padang Alan forest

An interior forest with a relatively even canopy approximately 30– 40 m tall and an understory dominated by small or prostrate shrubs, including *Euthemis minor* and *Ficus deltoidea*. Characteristic tree species include *Shorea albida*, *Calophyllum obliquinervum*, *Cratoxylum glaucum* and *Combretocarpus rotundatus*.

Samples were collected from four small ponds within each of the five vegetation– hydrological zones, yielding a total of 20 samples (Fig. 1). In each pond, compact sediment samples were collected from the center of the water body using a soil corer with a diameter of 2 cm at a depth of 15 cm. The sediment samples were immediately place into 15 ml centrifuge tubes. Due to shallow water tables (there were no sediment) at the Alan Bunga and Padang Alan forest sites, peat samples were collected and placed into resealable plastic bags. All samples were immediately transported to the lab in Sarawak Tropical Peat Research Institute. Cells in sediment and peat samples were filtered and concentrated following the protocol described by Lentendu et al. (2023). The sediment-filtered extract was processed for environmental DNA (eDNA) extraction within 24 hours.

### Environmental parameters and soil physicochemical properties

Soil water pH and conductivity were measured *in situ* using 913 Meter (Metrohm AG, Switzerland) from sample tubes. Soil temperature was measured using a portable soil temperature meter Hanna CheckTemp 1 (Hanna Instruments Inc, USA). Salinity was calculated from conductivity and temperature measurements. These environmental measurements were measured from four water ponds and mean values were used in subsequent analyses (Table S1). Water table readings were not taken during this study. However, average water table measurements for October 2023 were made available from Sarawak Tropical Peat Research Institute’s permanent water table pipes installed at each vegetation–hydrological zone except for coastal vegetation. The water table measurements were conducted as described in Dom et al. (2021). Additional peat physicochemical data, including humification index, loss on ignition, bulk density, carbon and nitrogen contents and C:N, were obtained from a separate, unpublished measurements and are reported in Table S2. The pyrophosphate solubility index (PSI) was determined according to the method by Karl (1956), a colorimetric estimation of the degree of humification of organic soil using an ultraviolet visible spectrometer UV/VIS Lambda 25 (Perkin Elmer, USA). Loss on ignition (LOI) was determined by weighing approximately 2.0 g of the peat soil sample and analyzed using thermo gravimetric analyzer TGA 701 (Leco, USA). Bulk density was assessed by collecting soil core samples using a core ring (5 cm diameter, 5.1 cm height). Total nitrogen (N) and total carbon (C) contents were determined using a TruMac CN Analyzer (LECO, USA), and the C/N ratio was subsequently calculated from the measured values. Available phosphorus was measured using Inductively Coupled Plasma-Optical Emission Spectrometry (ICP-OES) Agilent 5800 (Agilent Technologies, USA).

### eDNA extraction, PCR and sequencing

Approximately 0.25 g of sediment-filtered extract was used for eDNA extraction with the DNeasy PowerSoil Pro Kit (QIAGEN, Germany), following the manufacturer’s protocol. DNA purity and concentration were assessed using a NanoDrop spectrophotometer (Thermo Fisher Scientific, USA). Extracted eDNA was freeze and dried for downstream analyses.

The eDNA amplification targeting Arcellinida mitochondrial cytochrome oxidase subunit I (COI) gene was performed following the protocol described in González-Miguéns et al. (2023), available at dx.doi.org/https://doi.org/10.17504/protocols.io.yxmvm2389g3p/v1. We performed the first PCR with universal COI primers (Folmer et al., 1994), and the second PCR with a specific primer for Arcellinida (González-Miguéns et al., 2022) with a unique combination of tags for each sample to identify them. Amplified products were verified and purified by gel band excision. Purified DNA was quantified using an Invitrogen Qubit 3 fluorometer with the Thermo Fisher Scientific dsDNA High Sensitivity assay kit. Finally, purified DNA products were pooled and sequenced on an Illumina NextSeq 2000 platform using 600 cycles at the Genomic Unit of the Fundación Parque Científico de Madrid (Spain).

### eDNA Data curation and “tag jumping”

Illumina sequencing reads were processed according to the pipeline described in earlier work by González-Miguéns et al. (2023): (1) trimming of primers and demultiplexing was done with Cutadapt ver. 2.8 (Martin, 2011). (2) The resulting reads per sample were analyzed with the DADA2 R package (Callahan et al., 2016), filtering and trimming after positions 210 and 200 for the forward and reverse respectively. We dereplicated these sequences, merged the paired reads to generate amplicon sequencing variants (ASVs), and removed the chimeric sequences using the options proposed in DADA2 (Table S3). (3) For taxonomic assignation we applied VSEARCH ver. 2.14 (Rognes et al., 2016), using as a reference the eKOI database from González-Miguéns et al. (2024).

Samples were organized bioinformatically with the help of indexes unique to each sample, referred to as “tags,” which introduces the possibility of incorrect sample assignment, known as “tag jumping” (Rodriguez-Martinez et al., 2023; Schnell et al., 2015). We carried out a combination of protocols developed in earlier studies to reduce this bias (ElKhouri-Vidarte et al., 2025; González-Miguéns et al., 2023; Useros et al., 2024). According to the procedure outlined in (ElKhouri-Vidarte et al., 2025), the criteria required to eliminate incorrectly assigned sequences were established. Briefly, this method was based on the observation of tag combinations that do not match to any sample, and are therefore necessarily produced by tag jumping. The proportion of these “false reads” was used to set a threshold. First, we eliminated all ASVs below a minimum read count of 10 thus excluding those with too few reads that might be associated with errors or noise. Thereafter, we calculated the thresholds based on read proportions between wrong tag combinations and “real” samples. Finally, data were filtered to retain only ASVs assigned to the order Arcellinida with a percentage identity ≥84%. This number was determined empirically by González-Miguéns et al. (2024).

### Molecular OTU delineation and Phylogenetic Analysis

The ASVs assigned to Arcellinida were aligned with all known Arcellinida species for which DNA barcode sequences are available, as well as with the metabarcoding database of González-Miguéns et al. (2025), using the MAFFT auto algorithm (Katoh et al., 2002) implemented in Geneious version 2019.0.4. Finally, a distance matrix was generated to cluster the ASVs in operational taxonomic units (OTUs) using the R package DECIPHER ver. 2.22 (Wright, 2016). In order to approximate as much as possible our diversity inventory to a survey based on real species composition, we followed a barcoding gap which corresponds to the genetic distance between the sister species *Arcella salobris* and *Arcella uspiensis* (Useros et al., 2023). This barcoding gap corresponds to a distance of 2%. Based on this distance, we grouped the obtained ASVs (considered as haplotypes) into OTUs, the equivalent to species. Spearman’s rank correlation was used to assess the association between sequencing depth and OTU richness across samples.

Next, we proceeded to build de OTUs tree. Topologies and node supports were evaluated with maximum likelihood (ML), using IQTREE2 ver. 2.4 (Minh et al., 2020). The best substitution models were selected with ModelFinder (Kalyaanamoorthy et al., 2017), implemented in IQ-TREE2 ver. 2.4. Node supports were assessed with 10000 ultrafast bootstrap replicate approximation. Finally, the tree was built and represented with OTUs and barcode sequences of Arcellinida. The resulting tree was edited using FigTree v1.4.3 (Rambaut, 2016) and Affinity v3.2.0 (Canva, 2026).

### Diversity analysis

We generated a boxplot to compare OTU richness using the ggplot2 package (Wickham, 2016), and we generated one-way ANOVA using the aov() function from the stats package (Chambers et al., 1992) to assess those differences. Tukey’s HSD test was used for post-hoc pairwise comparisons. To visualize shared OTUs among vegetation–hydrological zones, we employed the ComplexUpset package (Krassowski et al., 2022) to create an UpSet diagram. We also created bar plots to compare the relative abundance of Arcellinida infraorders across vegetation–hydrological zones using the ggplot2 package. Then, we performed a non-metric multidimensional scaling (NMDS) to illustrate beta diversity patterns among communities associated with a pH gradient, using the function metaMDS() from vegan package (Oksanen et al., 2026). The analysis was based on presence–absence Jaccard distances and the resulting ordination was visualized using the ggplot2 package. We assessed the differences in community composition using permutational multivariate analysis of variance (PERMANOVA) with 999 permutations, using the adonis2() function from the vegan package.

To compare tropical peatlands with temperate and boreal peatlands, we generated bar plots using our relative abundance data (Table S4) and the “*Sphagnum* and wet mosses” data from García Bodelón et al. (2026), using the ggplot2 package.

All analysis were carried out using R software v4.5.1. The corresponding figures were prepared using Affinity v3.2.0 (Canva, 2026).

## RESULTS

### Environmental characteristics along the peat-dome transect

Environmental conditions varied along the peat-dome transect from the estuarine bank to the interior Padang Alan forest (Table S1). Among the measured variables, pH and water-table depth showed the clearest differences among vegetation–hydrological zones. The estuarine bank was distinguished by a comparatively higher pH (6.08 ± 0.098; range: 6.00 – 6.22), whereas the four inland zones were consistently acidic, with mean pH values ranging from 3.62 to 4.72. Water-table depth also varied across the peat-dome transect, ranging from –5.9 cm to –37.4 cm (Table S1). As for the river bank of Batang Lupar River, at the time of sampling, it was at low tide estimated at 1.82 meters below Mean Sea Level (MSL).

Peat physicochemical properties also differed among the four inland sites (Table S2). The pyrophosphate solubility index (PSI) was markedly higher in Mixed Peat Swamp forest hydrological zone (55.65) than in the other zones (4.31 – 6.50). The LOI analysis showed that the organic matter contents in the soil samples were more than 98% and ash content between 0.86 % to 1.39 %, which were of normal range for organic soil. The bulk density at the study sites was between 0.12 g cm^-3^ to 0.16 g cm^-3^. Alan Bunga also had the highest reported C:N (44.3), while available phosphorus values were lower in Alan Bunga and Padang Alan than in Mixed Peat Swamp and Alan Batu zones.

### Arcellinida taxonomic composition and phylogeny

The number of Illumina reads per sample through dada2 pipeline is detailed in Table S3. A total of 581 Arcellinida ASVs were identified in this study, none of which had been detected in any previous study (Fig. 2 and 3). After clustering ASVs at a 2% threshold, we obtained 225 OTUs in total, ranging between 1 and 41 per sample. Observed molecular OTU richness was not significantly related to the number of retained reads (ρ = 0.359, P = 0.120), which means that diversity has been satisfactorily sampled and therefore rarefaction or normalization was not deemed necessary to compare diversity between samples.

**Figure 2.**
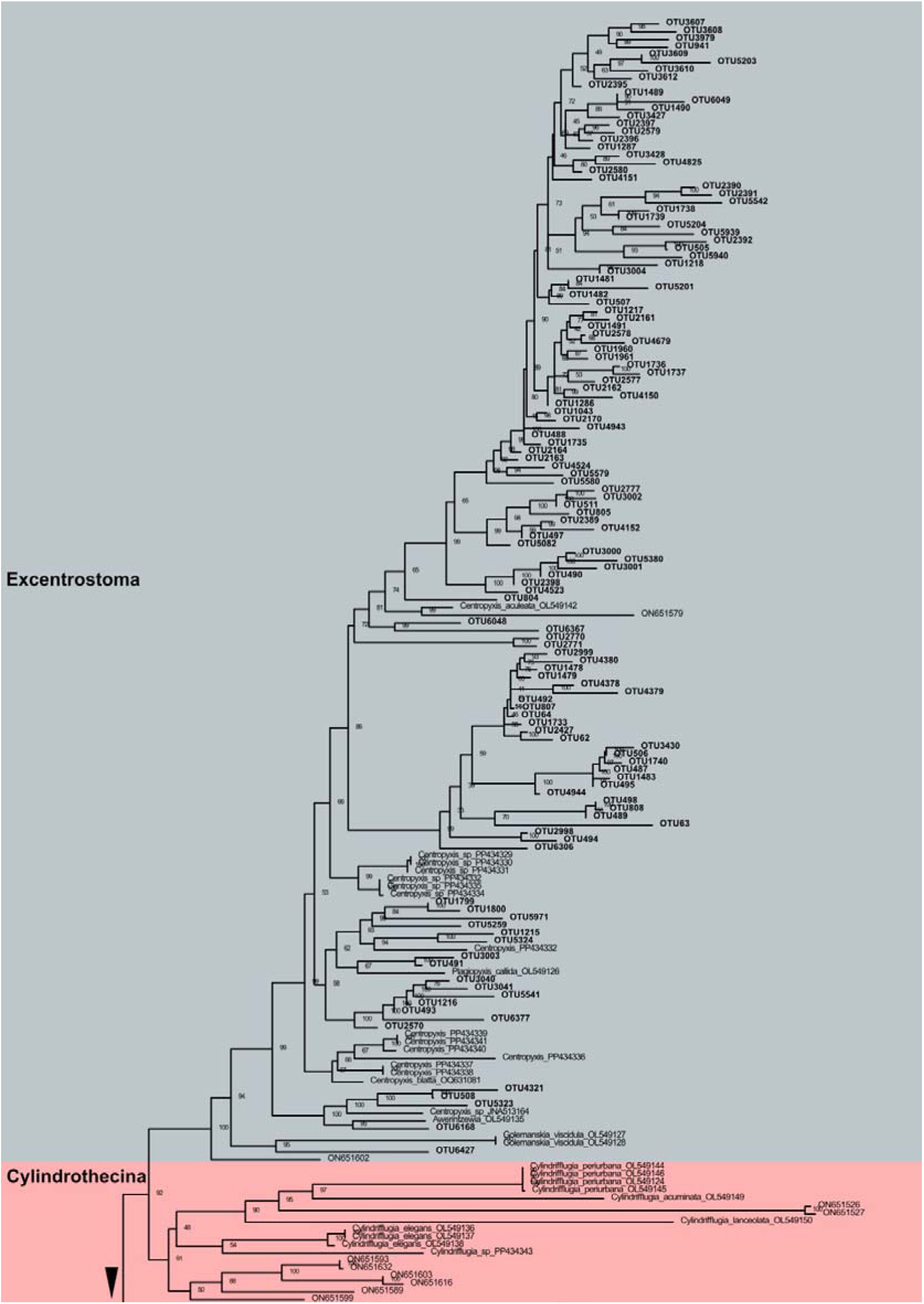
First section of the COI phylogenetic tree of Arcellinida OTUs performed using of the maximum-likelihood. New sequences found are designated as OTU in bold. Colors indicate infraorders, and bootstrap support values are shown at the corresponding nodes.

**Figure 3.**
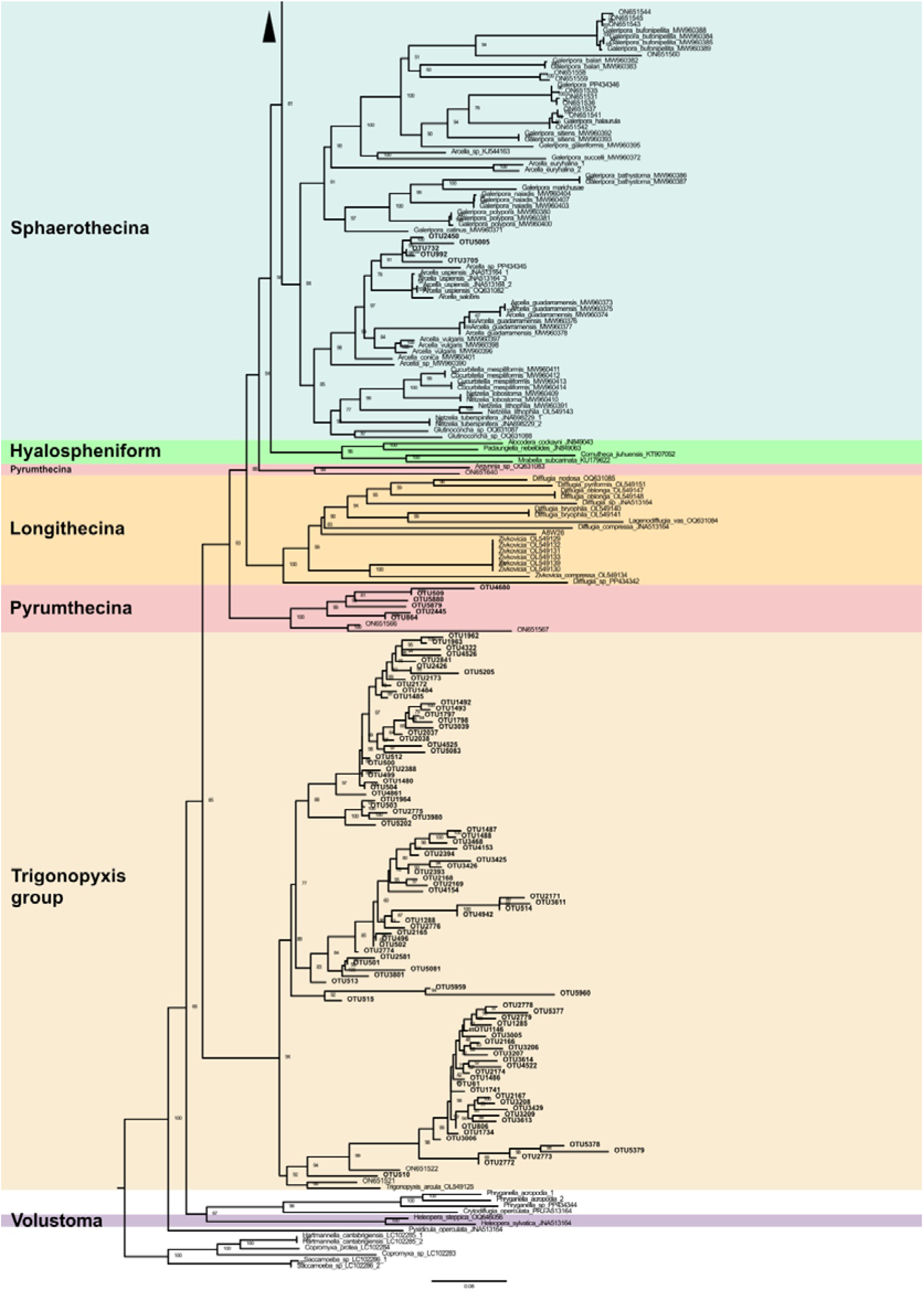
Second section of the COI phylogenetic tree of Arcellinida OTUs performed using of the maximum-likelihood. New sequences found are designated as OTU in bold. Colors indicate infraorders, and bootstrap support values are shown at the corresponding nodes.

Taxonomic composition of Arcellinida across all vegetation–hydrological zones, as revealed by phylogeny was the following: Infraorder Excentrostoma (125 OTUs), sequences related to genus *Trigonopyxis* (86 OTUs), infraorder Sphaerothecina (8 OTUs) and infraorder Pyrumthecina (6 OTUs) (Fig. 2 and 3). Notably, all *Trigonopyxis*-related OTUs clustered into a single, well-supported clade with 100% bootstrap support in the phylogenetic tree (Fig. 3).

We also compared our high-ranking taxa composition with similarly metabarcoded communities from six different temperate peatlands from Bulgaria (García-Bodelón et al., 2026) (Fig. S1). Our results differed notably as we have (1) a higher proportion of *Trigonopyxis-*related sequences, (2) a lower proportion of Pyrumthecina (3) and the absence of Hyalospheniformes related taxa group (Fig. S1). Moreover, we detected OTUs that branched with the sequences ON651566 and ON651567 reported by González-Miguéns et al. (2023). In Ribeiro et al. (2023) these sequences branched robustly with *Argynnia* sp. (OQ631083), but our phylogeny did not recover that clade; instead, they appeared split up into two separated clades (Fig. 3). However, we continue naming both clades as Pyrumthecina.

### Arcellinida richness and distribution among vegetation–hydrological zones

We found that OTU richness varied significantly according to vegetation–hydrological zones (ANOVA: F = 4.78, p = 0.011). The Alan Bunga forest showed significantly higher richness values than the areas of Coastal vegetation, Mixed Peat Swamp and Alan Batu forest (Fig. 4A). Richness values were comparatively similar among the four river-bank samples, whereas Padang Alan forest exhibited the greatest within-zone variability, ranging from 4 to 33 OTUs per sample (Fig. 4A).

**Figure 4.**
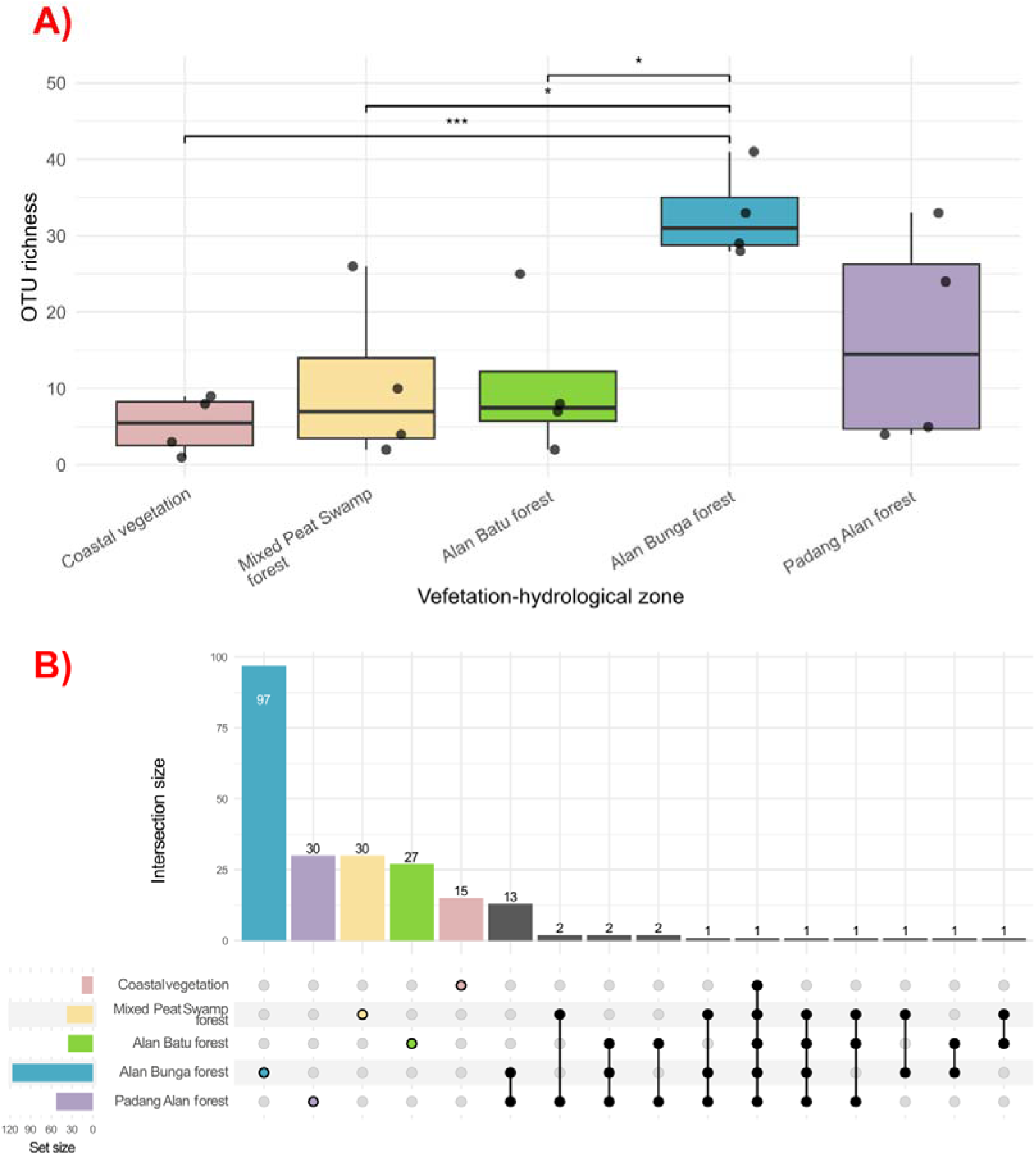
A) Boxplots represent the distribution of OTU richness values within each vegetation-hydrological zone. One-way ANOVA revealed significant differences among zones (p = 0.011), followed by Tukey’s HSD post-hoc test for pairwise comparisons. Significance codes are shown only for statistically significant pairwise comparisons: *** p < 0.001; ** p < 0.01; * p < 0.05. B) UpSet plot showing the distribution of OTUs among vegetation–hydrological zones. The upper bars indicate the number of OTUs that are unique to or shared among the combinations of vegetation–hydrological zones represented by the connected dots in the matrix below. The bars on the left-down show the total number of OTUs detected in each vegetation-hydrological zone.

Most OTUs exhibited restricted distributions along the transect (Fig. 4B). From a total of 225 OTUs detected, 199 OTUs (88.4%) were restricted to a single vegetation–hydrological zone, whereas only 26 (11.6%) occurred across two or more zones. Alan Bunga forest harbored the highest number of zone-restricted OTUs, with 97 OTUs found exclusively within this habitat (Fig. 4B). Only one OTU (OTU492) was shared across all five zones. Phylogenetic analysis placed OTU492 within the infraorder Excentrostoma (Fig. 2).

### Community composition of vegetation-hydrological zones

PERMANOVA revealed significant differences in community composition (F = 1.49, R² = 0.285, p = 0.001). Coastal vegetation zone was dominated by Excentrostoma (56.25%), followed by Sphaerothecina (37.5%) and a single OTU from the *Trigonopyxis* group. Mixed Peat Swamp and Alan Batu forest were both primary composed by the *Trigonopyxis* group (68.42% and 66.67% respectively), followed by Excentrostoma (23.68% and 27.78%, respectively), and a few Pyrumthecina. On the contrary, Alan Bunga forest and Padang Alan forest were dominated by Excentrostoma (68.38% and 81.13% respectively), followed by the *Trigonopyxis* group (28.21% and 15.09% respectively) and some Pyrumthecina. In addition, the Alan Bunga forest contained two Sphaerothecina OTUs (Fig. 5). NMDS analysis showed that communities from Coastal vegetation zone differed most from the rest of peatland communities. Furthermore, the samples from the Alan Bunga and Padang Alan forests were closely positioned in NMDS space (Fig. 6).

**Figure 5.**
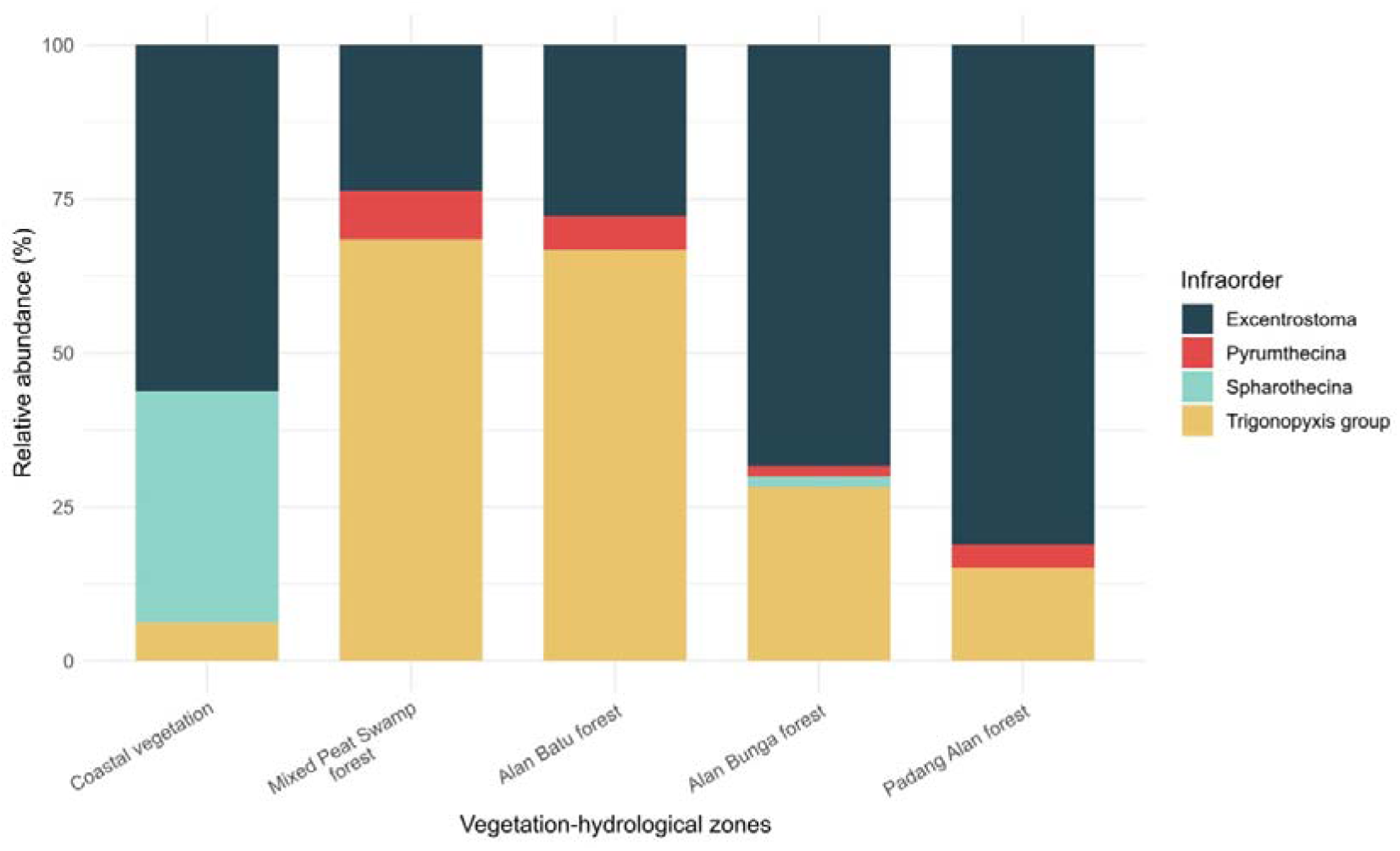
Relative abundance of Arcellinida infraorders across vegetation–hydrological zones. Bars represent vegetation–hydrological zones and colors indicate the proportion of OTUs assigned to each infraorder as identified in the legend.

**Figure 6.**
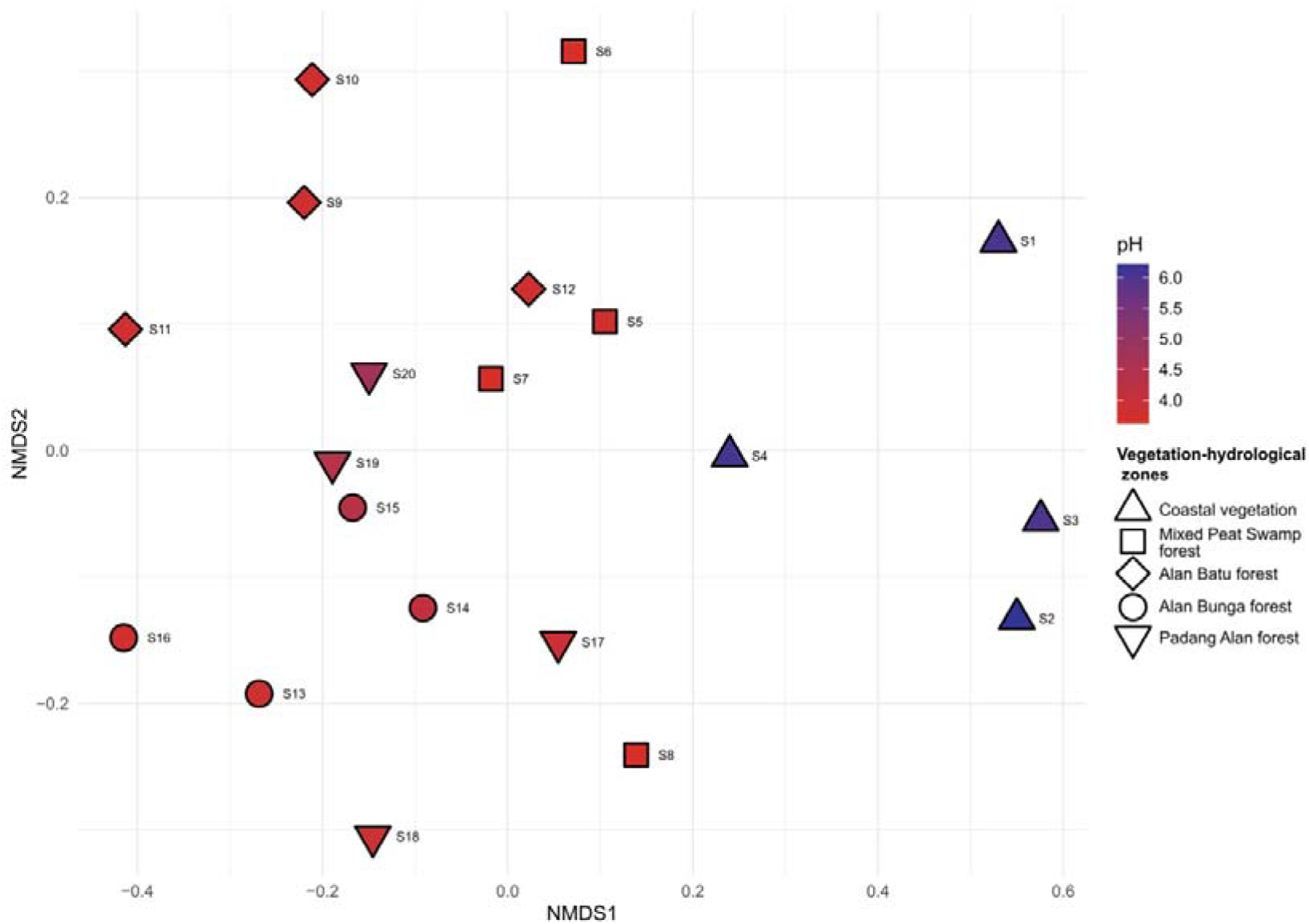
NMDS of the samples based on Jaccard dissimilarity (Stress = 0.091). Symbol shape indicates the vegetation–hydrological zones, and symbol fill color corresponds to pH values as shown in the legend.

## DISCUSSION

### Diversity and Composition of Arcellinida in Tropical Peatlands

This study identified new genetic lineages of Arcellinida inhabiting the tropical peat swamp forests of Maludam National Park. Indeed, none of the ASVs nor the OTUs detected in the study had been previously reported in any molecular survey (e.g. García-Bodelón et al., 2025; González-Miguéns et al., 2026; Soler-Zamora et al., 2026; Useros et al., 2025a). Although the novelty of molecular Arcellinida data from terrestrial South Asian environments may partly explain the distinctiveness of these communities, the marked differences observed even at high taxonomic levels suggest that the environmental conditions *per se* are a stronger driver of community composition.

The most notable feature of these communities is the prevalence of *Trigonopyxis*-related sequences. Such organisms have been abundantly reported in tropical ecosystems from South America and Southeast Asia based on microscopical observations (Krashevska et al., 2007, 2010; Tsyganov, Malysheva, et al., 2022). In the Ecuadorian Andes, Schulz et al. (2018) found several morphotypes possibly assignable to species, but as most individuals observed belonged to a morphological continuum, authors adopted a conservative approach. Here, we found no less than 86 OTUs, which formed a robust clade together with the voucher sequence *Trigonopyxis arcula* OL549125 (bootstrap: 100%) in the Arcellinida tree. This clade is rare in extratropical ecosystems; only two OTUs from this group were found in a *Sphagnum* patch in northern Spain (González-Miguéns et al., 2023). The genetic diversity found in the Maludam National Park suggests that *Trigonopyxis* and related organisms might form an independent infraorder. The huge molecular diversity can be also associated to the morphological diversity found in (Schulz et al., 2018). However, further sequencing of *Trigonopyxis* resemble organisms from different habitats, based on barcoded analyses, are needed in order to resolve their position in the Arcellinida tree and shed light on their systematics.

Our survey also revealed a significant diversity (8 OTUs) of organisms related to Pyrumthecina, also present in Northern Hemisphere peatlands (Bulgaria; García-Bodelón et al., 2026). This recently described infraorder (Porfirio-Sousa et al., 2025) appeared divided into two major subclades; while the Bulgarian sequences were related to *Argynnia* sp. (OQ631083) from freshwater (Ribeiro et al., 2023), our tropical peatland Pyrumthecina branched with sequences ON651566 and ON651567 reported by González-Miguéns et al., (2023) (Fig. S2) and originated from peatlands in France. A better taxonomic assessment of this infraorder will allow determining the ecological preferences in this group of mostly peatland associated amoebae.

The Arcellinida diversity revealed here contrasted strongly with that reported from temperate *Sphagnum*-dominated peatlands, such as those described in Bulgaria (Fig. S1; García-Bodelón et al., 2026). Total richness was comparable to boreal and temperate peatlands, that is, in the lower end of environmental diversity found in different temperate (García-Bodelón et al., 2026) and Mediterranean biota (ElKhouri-Vidarte et al., 2025). The composition, however, changed drastically. Beyond the *Trigonopyxis* group explained before, the recently described Pyrumthecina infraorder (Porfirio-Sousa et al., 2025) much more diversified and abundant in temperate peatlands; Hyalospheniformes, an abundant and well-studied group in European peatlands (Heger et al., 2013; Singer et al., 2019), were absent from Maludam samples (Fig. S1).

### Community turnover along the estuary-to-peat-dome gradient

The quality and degree of decomposition of peat vary depending on the vegetation– hydrological zones, ranging from mineral substrate in the coastal vegetation to highly decomposed peat in the Peat Swamp forest and less decomposed in Alan Batu/Bunga and Padang Alan forests (Melling, 2016). Accordingly, Dom et al. (2021) have shown a correlation between prokaryotic microbiomes and vegetation–hydrological zones in Sarawak peatlands. Our results are globally in line with these findings. The low number of OTUs shared across the complete transect (Fig. 4B) indicated pronounced spatial turnover in Arcellinida community composition (Fig. 4A).

Our physicochemical measurements provide additional context for the community turnover observed among hydrological zones. The substantially higher PSI values in mixed peat swamp forest indicate greater peat humification than at the other inland sites (Table S2), consistent with the pattern reported by Dom et al. (2021). On the other hand, Alan Bunga and Padang Alan zones, which supported together the highest OTU richness, had faster organic matter turnover; their highest C:N ratios and lower available phosphorous content indicate faster decomposition activity. Fast nutrient turnover in soil has been correlated with high diversity in another group of testate amoebae, the Euglyphida (Lara et al., 2016).

Alongside these differences in peat properties, vegetation structure may also help explain the observed patterns of Arcellinida richness. Alan Bunga and Padang Alan forests both have a relatively dense understory, which may increase resource availability and microhabitat heterogeneity. A positive relationship between plant and testate amoebae diversity have been shown in boreal and temperate peatlands, as well as in other temperate climate ecosystems (Tsyganov, Bobrov, et al., 2022; Tsyganov et al., 2023). Dense vegetation can provide fresh organic matter and macroaggregates that constitute the primary substrates for decomposers and bring additional microheterogeneity. Accordingly, both vegetation-hydrological zones hosted the highest prokaryotic diversity (Dom et al., 2021), suggesting possible indirect effects where prey diversity may influence predator richness. Habitat complexity may provide an additional explanation for the higher Arcellinida richness observed in Alan Bunga and Padang Alan forests. De Araujo et al. (2018) suggested that protist richness increased in climax vegetation because of the higher complexity of the leaf litter (or, here, the less decomposed peat). Furthermore, the pneumatophores present in Alan Bunga forest (Busman, Melling, et al., 2023) increase microscale habitat complexity, promoting environmental heterogeneity which could also affect positively Arcellinida richness. Finally, as Arcellinida have feeding preferences (Berlinches de Gea et al., 2024), differences in prokaryotic diversity may also influence Arcellinida diversity through bottom-up effects on the microbial food web.

The communities inhabiting in Coastal vegetation zone differ markedly from those found in other vegetation-hydrological zones, with notable differences also observed in soil pH (neutral versus very acidic) and the depth of the water table– factors that have been considered the main drivers of the composition of the amoeba communities studied in tropical peatlands (Krashevska et al., 2020; Swindles et al., 2014, 2018a). The influence of high tides could possibly be the strongest driver, as Arcellinida diversity has been found to decrease considerably in ecosystems under marine influence (Useros et al., 2025b).

## CONCLUSIONS

Altogether, tropical peatlands like the Maludam National Park in Malaysia host a unique Arcellinida diversity, which differs fundamentally from boreal and temperate peatlands or other temperate or Mediterranean ecosystems. While it is not especially species-rich, it is dominated by taxa that seldom appear in other biomes, including *Trigonopyxis*-related taxa, a particularly diverse clade. Furthermore, the different vegetation–hydrological zones host different diversity profiles, which suggest pronounced spatial turnover in Arcellinida community composition. These patterns were associated with concurrent changes in vegetation, pH, hydrology, substrate properties and tidal influence. Because these environmental characteristics covaried along a single transect, their individual contributions could not be separated conclusively. With these results, we aimed to provide a new framework to study tropical peatlands profile, and therefore promote the study and protection of these unique ecosystems.

## CREDIT AUTHORSHIP CONTRIBUTION STATEMENT

**Ángel García-Bodelón:** Conceptualization, Data curation, Formal analysis, Investigation, Methodology, Project administration, Visualization, Writing – original draft, Writing – review & editing. **Petr Heděnec**: Conceptualization, Funding acquisition, Project administration, Resources, Writing – review & editing. **Sharon Yu Ling Lau**: Conceptualization, Formal analysis, Validation of data, Funding acquisition, Writing – review & editing. **María Martínez-Ríos**: Data curation, Writing – review & editing. **Umar Hussaini Tarmizi**: Investigation, Methodology, Writing – review & editing. **Meldon Fienberg Kelbin:** Data curation, Formal analysis. **Kim San Lo**: Data curation, Formal analysis. Laurina Lenny Kambit: Data curation, Formal analysis. **Lulie Melling**: Funding acquisition, Writing – review & editing. **Nura ElKhouri-Vidarte**: Writing – review & editing. **Carmen Soler-Zamora**: Writing – review & editing. **Enrique Lara**: Conceptualization, Funding acquisition, Methodology, Project administration, Supervision, Validation, Writing – review & editing.

## DATA AVAILABILITY

All tables and figures, including the supplementary tables and figures associated with this study, are publicly available on Zenodo (https://doi.org/10.5281/zenodo.23213488).

## ACKNOWLEDGEMENTS

All soil and sediment samples were processed at the Molecular and Microbiology Research Division laboratories of the Sarawak Research and Development Council (former Sarawak Tropical Peat Research Institute).

## REFERENCES

1. Andriesse, J. P. (1988). Nature and Management of Tropical Peat Soils. Food & Agriculture Org.

2. Berlinches de Gea, A., Geisen, S., Grootjans, F., Wilschut, R. A., & Schwelm, A. (2024). Species-specific predation determines the feeding impacts of six soil protist species on bacterial and eukaryotic prey. European Journal of Protistology, 94, 126090. 10.1016/j.ejop.2024.126090

3. Birnbaum, C., Yusuf, A. Y., Yuwati, T. W., Rachmanadi, D., Kunarso, A., Bonner, M. T. L., & Grover, S. P. P. (2025). Soil Microbial Communities in Southeast Asian Peatlands: A Review. Mires and Peat, 32. 10.19189/001c.134066

4. Booth, R. K. (2008). Testate amoebae as proxies for mean annual water-table depth in Sphagnum-dominated peatlands of North America. Journal of Quaternary Science, 23(1), 43– 57. 10.1002/jqs.1114

5. Busman, N. A., Maie, N., Sangok, F. E., Melling, L., & Watanabe, A. (2023). Impacts of agricultural drainage on the quantity and quality of tropical peat soil organic matter in different types of forests. Geoderma, 439, 116670. 10.1016/j.geoderma.2023.116670

6. Busman, N. A., Melling, L., Goh, K. J., Imran, Y., Sangok, F. E., & Watanabe, A. (2023). Soil CO2 and CH4 fluxes from different forest types in tropical peat swamp forest. Science of The Total Environment, 858, 159973. 10.1016/j.scitotenv.2022.159973

7. Callahan, B. J., McMurdie, P. J., Rosen, M. J., Han, A. W., Johnson, A. J. A., & Holmes, S. P. (2016). DADA2: High-resolution sample inference from Illumina amplicon data. Nature Methods, 13(7), 581–583. 10.1038/nmeth.3869

8. Canva. (2026). Affinity (Version 3.2.0) [Computer software]. https://www.affinity.studio/es_es

9. Chambers, J. M., Freeny, A. E., & Heiberger, R. M. (1992). Analysis of Variance; Designed Experiments. In Statistical Models in S. Routledge.

10. Cheung, S. M. K., Pancost, R. D., Vreeken, M., Gallego-Sala, A. V., & Bryce, C. (2025). Structure and function of the global peatland microbiome (p. 2025.08.21.671631). bioRxiv. 10.1101/2025.08.21.671631

11. Creevy, A. L., Andersen, R., Rowson, J. G., & Payne, R. J. (2018). Testate amoebae as functionally significant bioindicators in forest-to-bog restoration. Ecological Indicators, 84, 274–282. 10.1016/j.ecolind.2017.08.062

12. Das, I., & Haas, A. (2010). New species of Microhyla from Sarawak: Old World’s smallest frogs crawl out of miniature pitcher plants on Borneo (Amphibia: Anura: Microhylidae). Zootaxa, 2571(1), 37–52. 10.11646/zootaxa.2571.1.2

13. de Araujo, A. S. F., Mendes, L. W., Lemos, L. N., Antunes, J. E. L., Beserra, J. E. A., de Lyra, M. Do C. C. P., Figueiredo, M. do V. B., Lopes, Â. C. de A., Gomes, R. L. F., Bezerra, W. M., Melo, V. M. M., de Araujo, F. F., & Geisen, S. (2018). Protist species richness and soil microbiome complexity increase towards climax vegetation in the Brazilian Cerrado. Communications Biology, 1(1), 135. 10.1038/s42003-018-0129-0

14. Dom, S. P., Ikenaga, M., Lau, S. Y. L., Radu, S., Midot, F., Yap, M. L., Chin, M.-Y., Lo, M. L., Jee, M. S., Maie, N., & Melling, L. (2021). Linking prokaryotic community composition to carbon biogeochemical cycling across a tropical peat dome in Sarawak, Malaysia. Scientific Reports, 11(1), 6416. 10.1038/s41598-021-81865-6

15. Dommain, R., Couwenberg, J., & Joosten, H. (2011). Development and carbon sequestration of tropical peat domes in south-east Asia: Links to post-glacial sea-level changes and Holocene climate variability. Quaternary Science Reviews, 30(7), 999–1010. 10.1016/j.quascirev.2011.01.018

16. ElKhouri-Vidarte, N., Useros, F., & Lara, E. (2025). Beneath the Cedars: Exploring the Water-Energy Balance on Arcellinida Biodiversity in Lebanon’s Cedar Forests. Microbial Ecology, 89(1), 33. 10.1007/s00248-025-02666-2

17. Folmer, O., Black, M., Hoeh, W., Lutz, R., & Vrijenhoek, R. (1994). DNA primers for amplification of mitochondrial cytochrome c oxidase subunit I from diverse metazoan invertebrates. 3(5), 294–299.

18. García-Bodelón, Á., Baković, N., Cano, E., Useros, F., Lara, E., & González-Miguéns, R. (2025). Predators in the Dark: Metabarcoding Reveals Arcellinida Communities Associated with Bat Guano, Endemic to Dinaric Karst in Croatia. Microbial Ecology, 87(1), 166. 10.1007/s00248-024-02483-z

19. García-Bodelón, Á., Todorov, M., ElKhouri-Vidarte, N., Soler-Zamora, C., Fournier, B., & Lara, E. (2026). Precipitation seasonality and soil carbon shape Arcellinida diversity across Bulgaria’s ecosystems (p. 2026.09.29.755277). bioRxiv. 10.64898/2026.09.29.755277

20. Gilbert, D., Amblard, C., Bourdier, G., & Francez, A.-J. (1998). The Microbial Loop at the Surface of a Peatland:Structure, Function, and Impact of Nutrient Input. Microbial Ecology, 35(1), 83–93. 10.1007/s002489900062

21. Gilbert, D., Mitchell, E. a. D., Amblard, C., Bourdier, G., & Francez, A.-J. (2003). Population dynamics and food preferences of the testate amoeba Nebela tincta major-bohemica-collaris complex (Protozoa) in a Sphagnum peatland. 42, 99–104.

22. González-Miguéns, R., Cano, E., García-Gallo Pinto, M., Peña, P. G., Rincón-Barrado, M., Iglesias, G., Blanco-Rotea, A., Carrasco-Braganza, M. I., de Salvador-Velasco, D., Guillén-Oterino, A., Tenorio-Rodríguez, D., Siemensma, F., Velázquez, D., & Lara, E. (2024). The voice of the little giants: Arcellinida testate amoebae in environmental DNA-based bioindication, from taxonomy free to haplotypic level. Molecular Ecology Resources, 24(7), e13999. 10.1111/1755-0998.13999

23. González-Miguéns, R., Cano, E., Guillén-Oterino, A., Quesada, A., Lahr, D. J. G., Tenorio-Rodríguez, D., de Salvador-Velasco, D., Velázquez, D., Carrasco-Braganza, M. I., Patterson, R. T., Lara, E., & Singer, D. (2023). A needle in a haystack: A new metabarcoding approach to survey diversity at the species level of Arcellinida (Amoebozoa: Tubulinea). Molecular Ecology Resources, 23(5), 1034–1049. 10.1111/1755-0998.13771

24. González-Miguéns, R., Cano, E., & Lara, E. (2025). Quaternary ice ages shaped protists phylogeography: The case of Arcellinida in the Iberian Peninsula (p. 2025.08.11.669755). bioRxiv. 10.1101/2025.08.11.669755

25. González-Miguéns, R., Cano, E., & Lara, E. (2026). Quaternary Ice Ages Shaped Protists Phylogeography: The Case of Arcellinida in the Iberian Peninsula. Molecular Ecology, 35(14), e70475. 10.1111/mec.70475

26. González-Miguéns, R., Gàlvez-Morante, A., Skamnelou, M., Antó, M., Casacuberta, E., Richter, D. J., Vaulot, D., Campo, J. del, & Ruiz-Trillo, I. (2024). A Novel Taxonomic Database for eukaryotic Mitochondrial Cytochrome Oxidase subunit I Gene (eKOI): Enhancing taxonomic resolution at community-level in metabarcoding analyses (p. 2024.12.05.626972). bioRxiv. 10.1101/2024.12.05.626972

27. González-Miguéns, R., Soler-Zamora, C., Villar-Depablo, M., Todorov, M., & Lara, E. (2022). Multiple convergences in the evolutionary history of the testate amoeba family Arcellidae (Amoebozoa: Arcellinida: Sphaerothecina): when the ecology rules the morphology. Zoological Journal of the Linnean Society, 194(4), 1044–1071. 10.1093/zoolinnean/zlab074

28. Heger, T. J., Mitchell, E. A. D., & Leander, B. S. (2013). Holarctic phylogeography of the testate amoeba Hyalosphenia papilio (Amoebozoa: Arcellinida) reveals extensive genetic diversity explained more by environment than dispersal limitation. Molecular Ecology, 22(20), 5172– 5184. 10.1111/mec.12449

29. Hodgkins, S. B., Richardson, C. J., Dommain, R., Wang, H., Glaser, P. H., Verbeke, B., Winkler, B. R., Cobb, A. R., Rich, V. I., Missilmani, M., Flanagan, N., Ho, M., Hoyt, A. M., Harvey, C. F., Vining, S. R., Hough, M. A., Moore, T. R., Richard, P. J. H., De La Cruz, F. B., … Chanton, J. P. (2018). Tropical peatland carbon storage linked to global latitudinal trends in peat recalcitrance. Nature Communications, 9(1), 3640. 10.1038/s41467-018-06050-2

30. IPBES. (2019). Global assessment report on biodiversity and ecosystem services of the Intergovernmental Science-Policy Platform on Biodiversity and Ecosystem Services. 10.5281/zenodo.6417333

31. Jassey, V. E. J., Signarbieux, C., Hättenschwiler, S., Bragazza, L., Buttler, A., Delarue, F., Fournier, B., Gilbert, D., Laggoun-Défarge, F., Lara, E., T. E. Mills, R., Mitchell, E. A. D., Payne, R. J., & Robroek, B. J. M. (2015). An unexpected role for mixotrophs in the response of peatland carbon cycling to climate warming. Scientific Reports, 5(1), 16931. 10.1038/srep16931

32. Kalyaanamoorthy, S., Minh, B. Q., Wong, T. K. F., von Haeseler, A., & Jermiin, L. S. (2017). ModelFinder: Fast model selection for accurate phylogenetic estimates. Nature Methods, 14(6), 587–589. 10.1038/nmeth.4285

33. Karl, A. Determination of the degree of humification in peat samples. (1956). J. Sci. Agric. Soc. Finland, 28, 18–35.

34. Katoh, K., Misawa, K., Kuma, K., & Miyata, T. (2002). MAFFT: A novel method for rapid multiple sequence alignment based on fast Fourier transform. Nucleic Acids Research, 30(14), 3059– 3066. 10.1093/nar/gkf436

35. Kottelat, M., Britz, R., Hui, T. H., & Witte, K.-E. (2006). Paedocypris, a new genus of Southeast Asian cyprinid fish with a remarkable sexual dimorphism, comprises the world’s smallest vertebrate. Proceedings of the Royal Society B: Biological Sciences, 273(1589), 895–899. 10.1098/rspb.2005.3419

36. Krashevska, V., Bonkowski, M., Maraun, M., & Scheu, S. (2007). Testate amoebae (protista) of an elevational gradient in the tropical mountain rain forest of Ecuador. Pedobiologia, 51(4), 319–331. 10.1016/j.pedobi.2007.05.005

37. Krashevska, V., Maraun, M., & Scheu, S. (2010). Micro-and macroscale changes in density and diversity of testate amoebae of tropical montane rain forests of Southern Ecuador. Acta Protozoologica, 49(1).

38. Krashevska, V., Tsyganov, A. N., Esaulov, A. S., Mazei, Y. A., Hapsari, K. A., Saad, A., Sabiham, S., Behling, H., & Biagioni, S. (2020). Testate Amoeba Species– and Trait-Based Transfer Functions for Reconstruction of Hydrological Regime in Tropical Peatland of Central Sumatra, Indonesia. Frontiers in Ecology and Evolution, 8. 10.3389/fevo.2020.00225

39. Krassowski, M., Arts, M., Lagger, C., & Max. (2022). krassowski/complex-upset: V1.3.5 [Computer software]. 10.5281/zenodo.7314197

40. Lara, E., Roussel-Delif, L., Fournier, B., Wilkinson, D.M. & Mitchell, E.A.D. (2016). Soil microorganisms behave like macroscopic organisms: patterns in the global distribution of soil euglyphid testate amoebae. J. Biogeogr., 43, 520–532. 10.1111/jbi.12660

41. Lamentowicz, M., Lamentowicz, Ł., & Payne, R. J. (2013). Towards quantitative reconstruction of peatland nutrient status from fens. The Holocene, 23(12), 1661–1665. 10.1177/0959683613508162

42. Lentendu, G., Bruni, E. P., Ah-Peng, C., Fujinuma, J., Kubota, Y., Lorite, J., Peñas, J., Huang, S., Strasberg, D., Vittoz, P., & Mitchell, E. A. D. (2023). Soil filtration-sedimentation improves shelled protist recovery in eukaryotic eDNA surveys. Molecular Ecology Resources, 23(6), 1361–1371. 10.1111/1755-0998.13808

43. Martin, M. (2011). Cutadapt removes adapter sequences from high-throughput sequencing reads. EMBnet.Journal, 17(1), 10–12. 10.14806/ej.17.1.200

44. Melling, L. (2016). Peatland in Malaysia. In M. Osaki & N. Tsuji (Eds), Tropical Peatland Ecosystems (pp. 59–73). Springer. 10.1007/978-4-431-55681-7_4

45. Minh, B. Q., Schmidt, H. A., Chernomor, O., Schrempf, D., Woodhams, M. D., von Haeseler, A., & Lanfear, R. (2020). IQ-TREE 2: New Models and Efficient Methods for Phylogenetic Inference in the Genomic Era. Molecular Biology and Evolution, 37(5), 1530–1534. 10.1093/molbev/msaa015

46. Morrison, E. S., Thomas, P., Ogram, A., Kahveci, T., Turner, B. L., & Chanton, J. P. (2021a). Characterization of Bacterial and Fungal Communities Reveals Novel Consortia in Tropical Oligotrophic Peatlands. Microbial Ecology, 82(1), 188–201. 10.1007/s00248-020-01483-z

47. Morrison, E. S., Thomas, P., Ogram, A., Kahveci, T., Turner, B. L., & Chanton, J. P. (2021b). Characterization of Bacterial and Fungal Communities Reveals Novel Consortia in Tropical Oligotrophic Peatlands. Microbial Ecology, 82(1), 188–201. 10.1007/s00248-020-01483-z

48. Oksanen, J., Simpson, G. L., Blanchet, F. G., Kindt, R., Legendre, P., Minchin, P. R., O’Hara, R. B., Solymos, P., Stevens, M. H. H., Szoecs, E., Wagner, H., Barbour, M., Bedward, M., Bengtsson, H., Bolker, B., Borcard, D., Borman, T., Carvalho, G., Chirico, M., … Weedon, J. (2026). vegan: Community Ecology Package (Version 2.7-5) [Computer software]. https://cran.r-project.org/web/packages/vegan/index.html

49. Page, S. E., Rieley, J. O., Shotyk, Ø. W., & Weiss, D. (1999). Interdependence of peat and vegetation in a tropical peat swamp forest. Philosophical Transactions of the Royal Society B: Biological Sciences, 354(1391), 1885–1897. 10.1098/rstb.1999.0529

50. Page, S. E., Rieley, J. O., & Wüst, R. A. J. (2006). Chapter 7 Lowland tropical peatlands of Southeast Asia. In Developments in Earth Surface Processes (Vol. 9, pp. 145–172). Elsevier. 10.1016/S0928-2025(06)09007-9

51. Porfirio-Sousa, A. L., Souza, M. B. G. E., Jones, R. E., Brown, M. W., Lahr, D. J. G., & Tice, A. K. (2025). Pyrumthecina Infraorder Nov.: Revisiting the morphological diversity and identifying the phylogenetic home of Argynnia within Arcellinida (Tubulinea:Amoebozoa) (p. 2025.08.20.671167). bioRxiv. 10.1101/2025.08.20.671167

52. Rambaut, A. (2016). FigTree (Version 1.4.3) [Computer software]. https://github.com/rambaut/figtree/releases/tag/1.4.3pre

53. Ribeiro, G. M., Useros, F., Dumack, K., González-Miguéns, R., Siemensma, F., Porfírio-Sousa, A. L., Soler-Zamora, C., Pedro Barbosa Alcino, J., Lahr, D. J. G., & Lara, E. (2023). Expansion of the cytochrome C oxidase subunit I database and description of four new lobose testate amoebae species (Amoebozoa; Arcellinida). European Journal of Protistology, 91, 126013. 10.1016/j.ejop.2023.126013

54. Ribeiro, K., Pacheco, F. S., Ferreira, J. W., de Sousa-Neto, E. R., Hastie, A., Krieger Filho, G. C., Alvalá, P. C., Forti, M. C., & Ometto, J. P. (2021). Tropical peatlands and their contribution to the global carbon cycle and climate change. Global Change Biology, 27(3), 489–505. 10.1111/gcb.15408

55. Rodriguez-Martinez, S., Klaminder, J., Morlock, M. A., Dalén, L., & Huang, D. Y.T. (2023). The topological nature of tag jumping in environmental DNA metabarcoding studies. Molecular Ecology Resources, 23(3), 621–631. 10.1111/1755-0998.13745

56. Rognes, T., Flouri, T., Nichols, B., Quince, C., & Mahé, F. (2016). VSEARCH: A versatile open source tool for metagenomics. PeerJ, 4, e2584. 10.7717/peerj.2584

57. Schnell, I. B., Bohmann, K., & Gilbert, M. T. P. (2015). Tag jumps illuminated – reducing sequence-to-sample misidentifications in metabarcoding studies. Molecular Ecology Resources, 15(6), 1289–1303. 10.1111/1755-0998.12402

58. Schulz, G., Maraun, M., Völcker, E., Scheu, S., & Krashevska, V. (2018). Evaluation of Morphological Characteristics to Delineate Taxa of the Genus *Trigonopyxis* (Amoebozoa, Arcellinida). Protist, 169(2), 190–205. 10.1016/j.protis.2018.02.005

59. Singer, D., Mitchell, E. A. D., Payne, R. J., Blandenier, Q., Duckert, C., Fernández, L. D., Fournier, B., Hernández, C. E., Granath, G., Rydin, H., Bragazza, L., Koronatova, N. G., Goia, I., Harris, L. I., Kajukało, K., Kosakyan, A., Lamentowicz, M., Kosykh, N. P., Vellak, K., & Lara, E. (2019). Dispersal limitations and historical factors determine the biogeography of specialized terrestrial protists. Molecular Ecology, 28(12), 3089–3100. 10.1111/mec.15117

60. Soler-Zamora, C., Cano, E., Vannucchi, P. E., Lara, E., & Fournier, B. (2026). Threshold responses of aquatic microbial diversity to terrestrial land use across Mediterranean catchments (p. 2026.08.07.743446). bioRxiv. 10.64898/2026.08.07.743446

61. Swindles, G. T., Baird, A. J., Kilbride, E., Low, R., & Lopez, O. (2018a). Testing the relationship between testate amoeba community composition and environmental variables in a coastal tropical peatland. Ecological Indicators, 91, 636–644. 10.1016/j.ecolind.2018.03.021

62. Swindles, G. T., Baird, A. J., Kilbride, E., Low, R., & Lopez, O. (2018b). Testing the relationship between testate amoeba community composition and environmental variables in a coastal tropical peatland. Ecological Indicators, 91, 636–644. 10.1016/j.ecolind.2018.03.021

63. Swindles, G. T., Reczuga, M., Lamentowicz, M., Raby, C. L., Turner, T. E., Charman, D. J., Gallego-Sala, A., Valderrama, E., Williams, C., Draper, F., Honorio Coronado, E. N., Roucoux, K. H., Baker, T., & Mullan, D. J. (2014). Ecology of Testate Amoebae in an Amazonian Peatland and Development of a Transfer Function for Palaeohydrological Reconstruction. Microbial Ecology, 68(2), 284–298. 10.1007/s00248-014-0378-5

64. Tsyganov, A. N., Bobrov, A. A., Shimano, S. D., Mitchell, E. A. D., Hagiwara, Y., Wall, A. A. J., Mazei, N. G., Chernyshov, V. A., Wanner, M., Zhong, Y., Sogame, Y., & Mazei, Y. A. (2022). Distribution of soil testate amoeba assemblages along an elevation gradient on Mount Fuji (Japan). European Journal of Protistology, 84, 125894. 10.1016/j.ejop.2022.125894

65. Tsyganov, A. N., Chertoprud, E. S., Mazei, N. G., Esaulov, A. S., Sadchikov, I. P., & Mazei, Y. A. (2023). The Effects of Vegetation and the Environment on Testate Amoeba Assemblages in Sphagnum Peatlands in the Northern Caucasus Mountains. Diversity, 15(2), 258. 10.3390/d15020258

66. Tsyganov, A. N., Malysheva, E. A., Mazei, Y. A., Anggi Hapsari, K., Behling, H., Sabiham, S., Biagioni, S., & Krashevska, V. (2022). Species– and Trait-Based Reconstructions of the Hydrological Regime in a Tropical Peatland (Central Sumatra, Indonesia) during the Holocene Using Testate Amoebae. Diversity, 14(12), 1058. 10.3390/d14121058

67. Useros, F., Fernández, L. D., González-Miguéns, R., Soler-Zamora, C., & Lara, E. (2025a). Inland Saline Lakes as Hotspots of Specialized, Nonmarine Protist Diversity: The Case of Arcellinida (Amoebozoa). Environmental DNA, 7(4), e70133. 10.1002/edn3.70133

68. Useros, F., Fernández, L. D., González-Miguéns, R., Soler-Zamora, C., & Lara, E. (2025b). Inland Saline Lakes as Hotspots of Specialized, Nonmarine Protist Diversity: The Case of Arcellinida (Amoebozoa). Environmental DNA, 7(4), e70133. 10.1002/edn3.70133

69. Useros, F., García-Cunchillos, I., Henry, N., Berney, C., & Lara, E. (2024). How good are global DNA-based environmental surveys for detecting all protist diversity? Arcellinida as an example of biased representation. Environmental Microbiology, 26(3), e16606. 10.1111/1462-2920.16606

70. Useros, F., González-Miguéns, R., Soler-Zamora, C., & Lara, E. (2023). When ecological transitions are not so infrequent: Independent colonizations of athalassohaline water bodies by Arcellidae (Arcellinida; Amoebozoa), with descriptions of four new species. FEMS Microbiology Ecology, 99(8), fiad076. 10.1093/femsec/fiad076

71. Wickham, H. (2016). Ggplot2. Springer International Publishing. 10.1007/978-3-319-24277-4

72. Wright, E. S. (2016). Using DECIPHER v2.0 to Analyze Big Biological Sequence Data in R. The R Journal, 8(1), 352–359. 10.32614/RJ-2016-025

73. Yu, Z. C. (2012). Northern peatland carbon stocks and dynamics: A review. Biogeosciences, 9(10), 4071–4085. 10.5194/bg-9-4071-2012

74. Zinck, J. A. (2011). Tropical and Subtropical Peats: An Overview. In J. A. Zinck & O. Huber (Eds), Peatlands of the Western Guayana Highlands, Venezuela: Properties and Paleogeographic Significance of Peats (pp. 5–28). Springer. 10.1007/978-3-642-20138-7_2

