## Supplementary material for "Tropical Peatlands as Hotspots of Hidden Arcellinida Diversity: Insights from Maludam National Park, Borneo (Sarawak, Malaysia)": Suplementary data

| Supplementary tables and figures | |
| --- | --- |
| Table S1 | Page 2 |
| Table S2 | Page 3 |
| Table S3 | Page 4 |
| Table S4 | Page 4 |
| Figure S1 | Page 5 |
| Figure S2 | Page 5 |

Table S1. Samples and measures variables.

| Sample | Forest type | Distance estimation from riverbank (m) | pH | Conductivity (uS/cm) | Salinity (mg/L) | Average water table depth |
| --- | --- | --- | --- | --- | --- | --- |
| RB-1 | Coastal vegetation | 50 | 6.03 | 5.09 | 3.2576 | ≈ 1.82 m below MSL |
| RB-2 | Coastal vegetation | 50 | 6.22 | 5.03 | 3.2192 |  |
| RB-3 | Coastal vegetation | 50 | 6 | 917.7 | 587.328 |  |
| RB-4 | Coastal vegetation | 50 | 6.07 | 478.6 | 306.304 |  |
| MA-1 | Mixed Peat Swamp | 1230 | 3.78 | 120.9 | 77.376 | -5.9 cm |
| MA-2 | Mixed Peat Swamp | 1230 | 3.64 | 119.1 | 76.224 |  |
| MA-3 | Mixed Peat Swamp | 1230 | 3.64 | 117.1 | 74.944 |  |
| MA-4 | Mixed Peat Swamp | 1230 | 3.62 | 116.2 | 74.368 |  |
| MB-1 | Alan Batu | 4460 | 3.79 | 87.2 | 55.808 | -25.4 cm |
| MB-2 | Alan Batu | 4460 | 3.79 | 86 | 55.04 |  |
| MB-3 | Alan Batu | 4460 | 3.79 | 85.1 | 54.464 |  |
| MB-4 | Alan Batu | 4460 | 3.79 | 88.07 | 56.3648 |  |
| MC-1 | Alan Bunga | 5900 | 3.82 | 80.6 | 51.584 | -37.4 cm |
| MC-2 | Alan Bunga | 5900 | 4.1 | 43 | 27.52 |  |
| MC-3 | Alan Bunga | 5900 | 4.3 | 13.4 | 8.576 |  |
| MC-4 | Alan Bunga | 5900 | 3.76 | 26.3 | 16.832 |  |
| MD-1 | Padang Alan forest | 8580 | 3.85 | 79.3 | 50.752 | -35.8 cm |
| MD-2 | Padang Alan forest | 8580 | 3.87 | 73.2 | 46.848 |  |
| MD-3 | Padang Alan forest | 8580 | 4.33 | 20.6 | 13.184 |  |
| MD-4 | Padang Alan forest | 8580 | 4.72 | 20.7 | 13.248 |  |

Table S2. Table S2. Physicochemical properties of soil at Mixed Peat Swamp forest (MPS), Alan Batu forest (ABt), Alan Bunga forest (ABg), and Padang Alan forest (PA) at 0–25 cm. Water table depth (WT) with (−) values indicating depths below soil surface. PSI: Pyrophosphate solubility index, LOI: Loss-on-ignition, BD: Bulk density, TC: Total carbon, TN: Total nitrogen, P: Phosphorus.

| **Site** | **Water Table Depth (cm)** | **Temp at 5cm (˚ C)** | **PSI** | **Ash (%)** | **LOI (%)** | **BD**  **(g cm^-3^)** | **TC (%)** | **TN (%)** | **C:N** | **Available P (ppm)** |
| --- | --- | --- | --- | --- | --- | --- | --- | --- | --- | --- |
| Mixed Peat Swamp (MPS) | -5.9 | 24.0 | 55.65 | 1.39 | 98.62 | 0.16 | 55.15 | 1.49 | 37.0 | 141.42 |
| Alan Batu (ABt) | -25.4 | 24.3 | 6.50 | 0.97 | 99.03 | 0.13 | 50.91 | 1.60 | 31.8 | 132.19 |
| Alan Bunga (ABg) | -37.4 | 24.0 | 4.31 | 0.86 | 99.14 | 0.13 | 50.02 | 1.13 | 44.3 | 81.76 |
| Padang Alan (PA) | -35.9 | 24.1 | 5.73 | 1.03 | 98.98 | 0.12 | 49.76 | 1.36 | 36.6 | 66.53 |

Table S3. Number of Illumina reads per sample through the DADA2 pipeline.

|  | input | filtered | denoisedF | denoisedR | merged | nonchim | Percentage of Arcellinida sequences |
| --- | --- | --- | --- | --- | --- | --- | --- |
| RB-1 | 412555 | 293426 | 291022 | 291630 | 278030 | 266270 | 64.5416975 |
| RB-2 | 835709 | 672667 | 666699 | 669534 | 648376 | 604454 | 72.3282865 |
| RB-3 | 723033 | 580165 | 575339 | 577147 | 560155 | 529062 | 73.1725938 |
| RB-4 | 561252 | 441449 | 436512 | 437820 | 420959 | 387124 | 68.9750771 |
| MA-1 | 485241 | 387753 | 385946 | 385490 | 365053 | 352990 | 72.7452956 |
| MA-2 | 703876 | 574266 | 571097 | 572886 | 547687 | 527212 | 74.901261 |
| MA-3 | 653268 | 538831 | 537754 | 537495 | 522521 | 511123 | 78.2409363 |
| MA-4 | 519686 | 422765 | 419852 | 421365 | 407100 | 384328 | 73.9538875 |
| MB-1 | 535969 | 444776 | 442974 | 443101 | 433201 | 412516 | 76.9663917 |
| MB-2 | 661395 | 531565 | 529347 | 529363 | 509594 | 482996 | 73.0268599 |
| MB-3 | 625089 | 517249 | 513847 | 514231 | 497765 | 472837 | 75.6431484 |
| MB-4 | 494909 | 411221 | 409623 | 410204 | 402433 | 387060 | 78.2083171 |
| MC-1 | 592031 | 494274 | 490783 | 491959 | 469918 | 452773 | 76.4779209 |
| MC-2 | 759479 | 612542 | 609081 | 610429 | 589981 | 572754 | 75.4140667 |
| MC-3 | 529304 | 442534 | 440834 | 440300 | 433957 | 389532 | 73.593247 |
| MC-4 | 615101 | 500479 | 497682 | 499436 | 445619 | 428179 | 69.6111695 |
| MD-1 | 481861 | 397507 | 395221 | 395524 | 389155 | 373156 | 77.4405897 |
| MD-2 | 750409 | 539333 | 535840 | 535993 | 520547 | 492246 | 65.5970278 |
| MD-3 | 718769 | 590299 | 587160 | 588511 | 574195 | 535836 | 74.549125 |
| MD-4 | 722538 | 603146 | 598855 | 601660 | 588455 | 571765 | 79.1328622 |

Table S4. This table is provided as a separate file in CSV format (Table S4.csv).

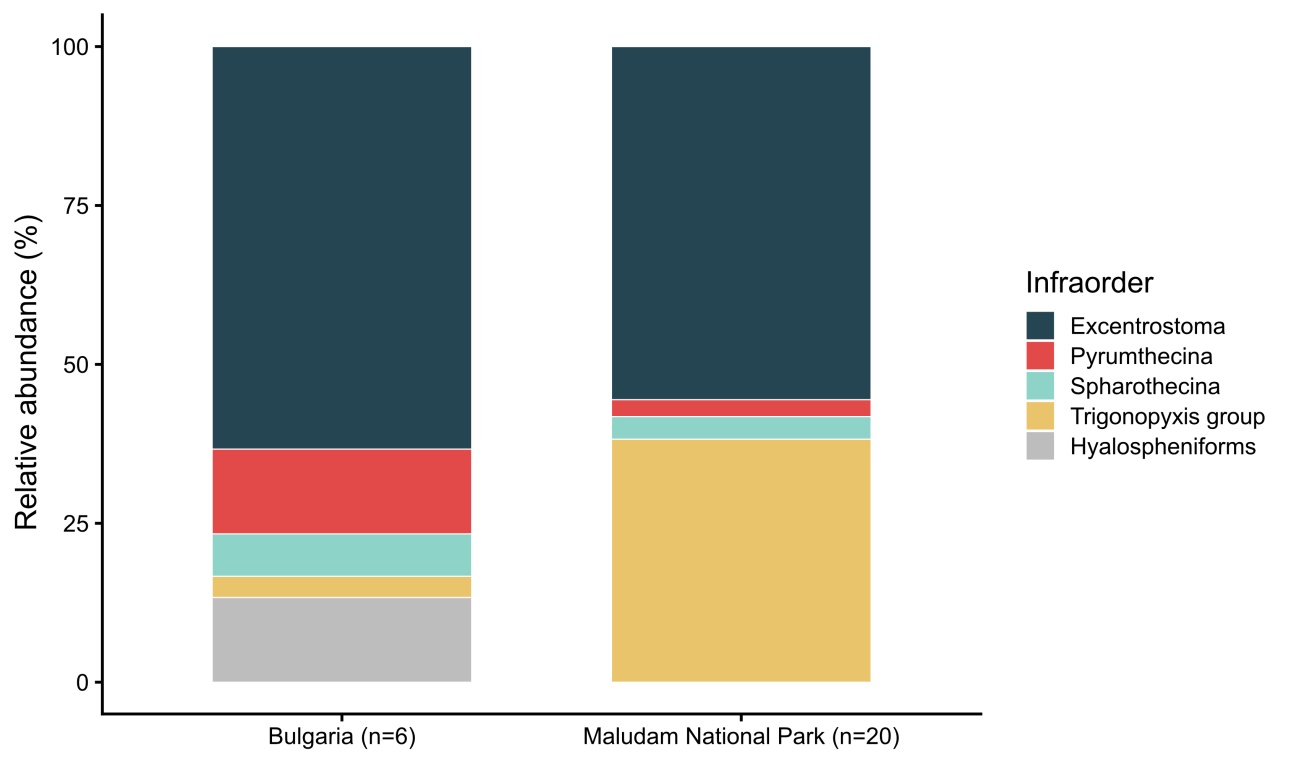

Figure S1. Infraorder composition of Arcellinida OTUs in temperate peatlands from Bulgaria and tropical peatlands from Maludam National Park (Malaysia).

Figure S2. Maximum-likelihood phylogenetic tree of Arcellinida OTUs based on mitochondrial cytochrome c oxidase subunit I (COI) sequences. Node support was assessed using the ultrafast bootstrap approximation with 10000 replicates. New sequences found are designated as OTU in bold. The dataset also includes OTUs recovered from *Sphagnum* and wet moss samples collected in temperate peatlands in Bulgaria (in red) (García-Bodelón et al., 2026.), together with reference sequences from barcoded species. This table is provided as a separate file in PDF format (Figure S2.pdf).
